# Are you suppressed? Viral load test coverage for people living with HIV in Mutare District, Manicaland Province, Zimbabwe, 2015-2017 Investigators

**DOI:** 10.1101/324350

**Authors:** Charles Uzande, Jeffery Edwards, Philip Owiti, Admire Tatenda Maravanyika, Simba Mashizha, Patron Titsha Mafaune

**Affiliations:** Provincial Medical Directorate (MOHCC) Manicaland Province Zimbabwe; Médecins Sans Frontières, Operational Centre Brussels, Brussels Belgium; Department of Global Health, University of Washington, Seattle, Washington, USA; International Union Against Tuberculosis and Lung Disease (The Union), Paris, France; National Tuberculosis, Leprosy and Lung Disease Programme, Nairobi, Kenya; District Medical Office (MOHCC) Mutare Manicaland Zimbabwe

## Abstract

**Background:** The third 90-90-90 UNAIDS goal require that 90% of people living with HIV (PLHIV) on antiretroviral treatment (ART) achieve viral load (VL) suppression. This study assessed the proportion of VL suppression and related factors among PLHIV on 1^st^ and 2^nd^ line ART in Mutare District, Manicaland Province, Zimbabwe between 2015-2017.

**Methods:** A retrospective study using routine HIV programme data from the electronic monitoring system for nine health facilities in Mutare District. VL suppression was defined as < 1,000 copies/ml.

**Results:** Of 16,590 registered patients, 15,566(94%) were on first-line and 1024(6%) on second-line ART. Of those on 1^st^-line ART, 2856(18%) had a VL test result documented, while 367(36%) of 2^nd^-line ART patients had VL results. VL suppression rates were 86% among those on 1^st^-line and 45% in 2^nd^-line ART. Independent risk factors associated with VL non-suppression for those on 1^st^-line ART were age 0-9 years (adjusted relative risk, aRR=2.9; 95% confidence interval, CI=1.7-4.8;P<0.001), 10-19 years (aRR=2.2;95%CI=1.4-3.2,P<0.001) compared to those 20-49 years, concurrent TB (aRR=9; CI=3.0-29.7,P<0.001) and male gender (aRR=1.5,95%CI=1.1-2.1;P=0.02). There were no significant risk factors associated with VL non-suppression for 2^nd^-line ART patients.

**Conclusion:** For PLHIV on 1^st^-line ART in Mutare district, Manicaland, Zimbabwe, the frequency of reported VL results were only 18% among those on 1^st^-line ART, while the rate of VL suppression was near 90%. Viral Load testing coverage appears to be lagging behind current Zimbabwe goals and increased support is needed to improve the quality of HIV care and help reduce the threat of possible HIV drug resistance in the future.

## Introduction

In December 2013, the Joint United Nations Programme on HIV/AIDS (UNAIDS) proposed ambitious new targets for HIV treatment scale-up beyond 2015. The new “90-90-90” goals for 2020 included 90% of all people living with HIV (PLHIV) know their HIV status, 90% diagnosed with HIV infection receive sustained antiretroviral therapy (ART) and 90% receiving ART will have durable viral load suppression.(1) UNAIDS data from 168 countries reveal progress and gaps across the HIV testing and treatment cascade with approximately 82% of those on treatment virally suppressed. Fully achieving the “90-90-90” targets translates to 73% of PLHIV virally suppressed; so far 7 countries including Botswana have already achieved or exceeded this target.(2)

In Zimbabwe, HIV prevalence has declined from 18% in 2009 to 14% in 2015(3) and there are an estimated 1.2 million PLHIV, with almost one million currently on ART. However, there remain challenges to achieving the projected 90-90-90 UNAIDS goals. In response, the Zimbabwe Ministry of Health and Child Care (MOHCC) has adopted viral load (VL) as the standard for ART monitoring, aiming to provide testing services to at least 90% of PLHIV receiving ART by 2018. To facilitate the national rollout, the Viral Load Scale Up Plan 2015-2018 was developed to assist in accelerating the scaling up of the VL testing through defined national testing targets and timeframes for achieving those targets.(4)

Manicaland province started rolling out VL monitoring in 2015 with support from partners. Currently all the seven districts within Manicaland are offering VL testing at selected high volume sites. Since 2015, Mutare Provincial Hospital (MPH) has been providing VL testing to all sites in the province. To date, no studies have been done in Manicaland to determine factors associated with non-suppression of VL.

Factors associated with VL non-suppression in other countries of sub-Saharan Africa (SSA) have included children and adolescents, not adhering to treatment, active TB, male gender, Who stage 4 and lower CD4 at ART initiation, while being on 2^nd^ or 3^rd^ line ART regimens were protective against non-suppression.(5)(6)(7)(8)

According to the Zimbabwe Population Based HIV Impact Assessment Survey 2015-2016 (ZIMPHIA), viral load suppression among adult HIV patients (15-64 years) on ART in Zimbabwe was 86.5%, however for Manicaland it was 61.5% for both PLHIV on ART and those not ART, the viral load suppression rates and non suppression for those on ART in the province was unknown.(9) Thus the aim of this study was to determine the proportion with non viral load suppression and related factors among PLHIV on 1^st^ and 2^nd^ line ART in Mutare District, Manicaland Province, Zimbabwe between 2015-2017.

## Materials and methods

**Design:** This was a retrospective study utilizing previously collected routine programmatic data from 2015-2017.

**Setting:** Zimbabwe is a low income country in southern Africa with a high burden of HIV. The estimated
population of Zimbabwe is 16 million and healthcare is largely provided free of cost by MOHCC supported facilities countrywide. Healthcare access remains a challenge, especially in rural areas, with a low physician and nurse to patient ratio of 1.6/10,000 and 7.2/10,000, respectively.(10) Despite healthcare access difficulties, with the support of multiple partners over the last decade, the cost of diagnosis and treatment with ART has been essentially cost free to everyone.

Mutare District is one of the seven districts in Manicaland province which is in south eastern Zimbabwe, with the provincial capital city, Mutare. The district (both urban and rural) has a population of 450,000.(11) All MOHCC facilities provide HIV care based upon the current Zimbabwe HIV guidelines which are consistent with the WHO 2015 recommendations.(12)

**Specific Setting:** There are 53 health facilities in Mutare district of which 48 of these are offering routine viral load testing. Ten of these sites, which are high volume, are supported by Médecins Sans Frontières (MSF).(13) The sites started targeted VL testing in September 2015 and by August 2016 routine VL testing was being done in 48 health facilities. For the ten high volume sites, HIV patient level data is captured in the MOHCC supported Electronic Patient Monitoring System (EPMS).

VL suppression is defined as having a VL of <1000 copies/ml of blood while non suppression is having a VL of >1000 copies/ml of blood. It is recommended that the first viral load test be taken at 6 months after ART initiation, 12 months and yearly. If non suppressed Extended Adherence Counselling (EAC) is done and a repeat test is done after 3 months, if it remains unsuppressed the patient is switched to second line ART according to the national guidelines.(14)

All VL tests for Manicaland and Masvingo Provinces are completed at MPH laboratory on a single MSF-supported VL platform. In 2017, MPH ran approximately 21,000 VL tests for both provinces. VL samples are transported from facilities outside of MPH by either whole blood, plasma (if centrifuge is available) in EDTA tubes requiring a cold-chain or by dried blood spot. VL sample transportation is specifically supported by collaborating partners including USAID and the Global Fund. The MPH reported sample rejection rate was approximately 2-3% in 2017. All MPH VL results are entered into the BIKA Laboratory Management Information System (LMIS).

**Study Participants:** The study population included all PLHIV in 9 HFs within Mutare District during the period from 2015-2017 who have been on either 1^st^ line or 2^nd^ line ART for at least 6 months. For ART patients with multiple VL tests, the most recent result was considered for analysis.

**Data Variables:** Data from nine health facilities (HFs) within Mutare District was extracted from the EPMS database into Excel (Microsoft, Redmond, WA, USA). Data variables included anonymised patient demographic and clinical characteristics, including viral load results.

**Statistical analysis:** Data variables were imported into EpiData Analysis software (*v2.2.2.182, EpiData Association, Odense, Denmark*) analysed and presented in frequencies for categorical data and continuous variables were categorised for further analysis. Proportion with/without viral load suppression was presented with 95% confidence intervals (CIs). Factors associated with non-suppression of viral load were assessed using chi-square test at bivariate level and log-binomial regression at multivariate level (using Stata/SE 14.1, *StataCorp, College Station, TX, USA*). Only variables significant at bivariate level and those which improved the fit of the models to the data were included in the final models. Relative Risks (RRs) were used to calculate the measures of association, and presented with their 95% CIs. Levels of significance were set at ≤ 5%.

## Ethics

The study was reviewed and approved by the Medical Research Council of Zimbabwe (Harare, Zimbabwe) and the International Union Against Tuberculosis and Lung Disease Ethics Advisory Group (Paris, France). Permission was granted by the Zimbabwe Ministry of Health and Child Care (Harare, Zimbabwe). There was no need for informed consent as we utilized anonymized data.

## Results

There were 16590 patients registered within the 9 Mutare District health facilities between January 2015 – December 2017, of whom 15,566 (94%) were on first line and 1,024 (6%) on second line ART regimens. Of those on 1^st^ line ART, 2856(18%) had a VL test results documented, while 367(36%) of 2^nd^ line ART patients had VL results, see (**Fig 1**). Those patients on 2^nd^ line therapy were significantly younger compared with those on a 1^st^ line regimen (38 years (interquartile range, IQR: 19-48) vs 43 years (IQR 35-50), *P<0.001*), more likely to be female (55% vs 33%, *P<0.001*) and single (30% vs 17%, *P<0.001*) see **Table 1**.

**Fig 1.**
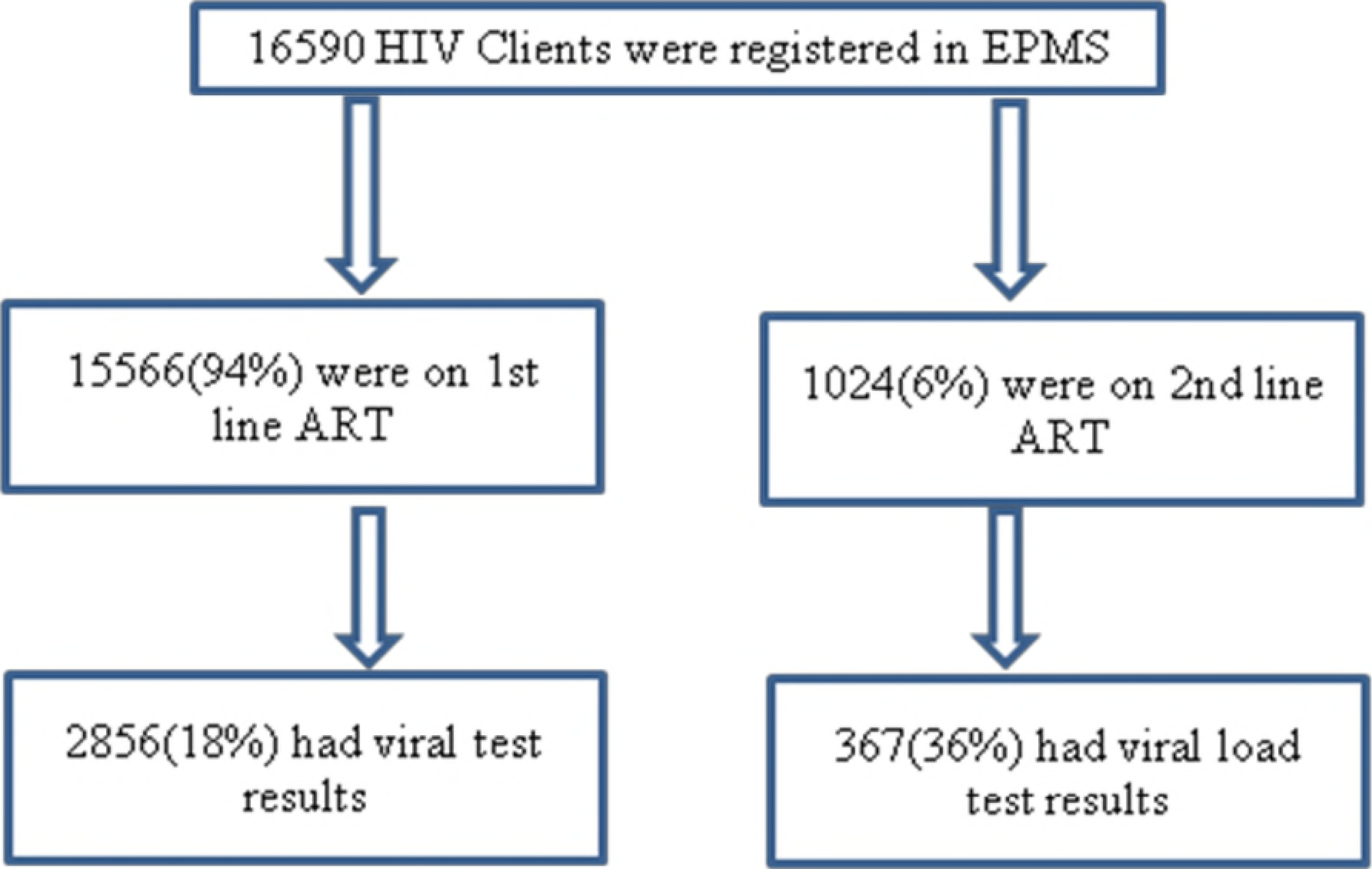
Numbers and proportions of HIV clients included and excluded at each stage for final analysis, Mutare District, Manicaland Province, Zimbabwe 2015-2017.

**Table 1.** *Demographic characteristics of HIV patients on 1^st^ and 2^nd^ line ART with viral load testing results in Mutare District, Manicaland Province, Zimbabwe 2015-2017 (N=3,223).*

| Variable | 1<sup>st</sup> line ART |  | 2<sup>nd</sup> line ART |  | Total | *p-value* |
| --- | --- | --- | --- | --- | --- | --- |
|  | N | (%) | N | (%) |  |  |
| **Age group (years)** |  |  |  |  |  |  |
| 0-9 | 58 | (2) | 16 | (4) | 76 |  |
| 10-19 | 137 | (5) | 82 | (3) | 229 |  |
| 20-49 | 1881 | (66) | 190 | (11) | 1929 |  |
| ≥50 | 780 | (27) | 79 | (15) | 845 |  |
| Total | 2856 | (100) | 367 | (100) | 3223 | *p<0.001* |
| Median age | 43 | IQR: 35-50 | 38 | IQR: 19-48 |  |  |
| **Gender** |  |  |  |  |  |  |
| Male | 929 | (33) | 167 | (46) | 1096 |  |
| Female | 1927 | (68) | 200 | (55) | 2127 |  |
| Total | 2856 | (100) | 367 | (100) | 3223 | *p<0.001* |
| **Marital Status** |  |  |  |  |  |  |
| Married | 1452 | (55) | 121 | (39) | 1573 |  |
| Divorced | 224 | (8) | 26 | (9) | 250 |  |
| Single | 400 | (15) | 107 | (35) | 507 |  |
| Widowed | 586 | (22) | 53 | (17) | 639 |  |
| Total | 2663 | (100) | 307 | (100) | 2969 | *p<0.001* |
| **Level of Education** |  |  |  |  |  |  |
| None | 35 | (3) | 7 | (5) | 42 |  |
| Primary | 318 | (26) | 43 | (30) | 361 |  |
| Secondary | 785 | (63) | 79 | (55) | 864 |  |
| Tertiary | 104 | (8) | 14 | (10) | 118 |  |
| Total | 1242 | (100) | 143 | (100) | 1385 | *p=0.2* |

IQR = interquartile range

Patients on 2^nd^ line ART were more likely to have presented with advanced WHO 3-4 stage disease than 1^st^ line (48% vs 42%, *P=0.03*), a VL < 1000 (45% vs 14%, *P<0.001*) and CD4 count < 100 (48% vs 25%,*p<0.001*), compared to 1^st^ line patients, see **Table 2.**

**Table 2.**
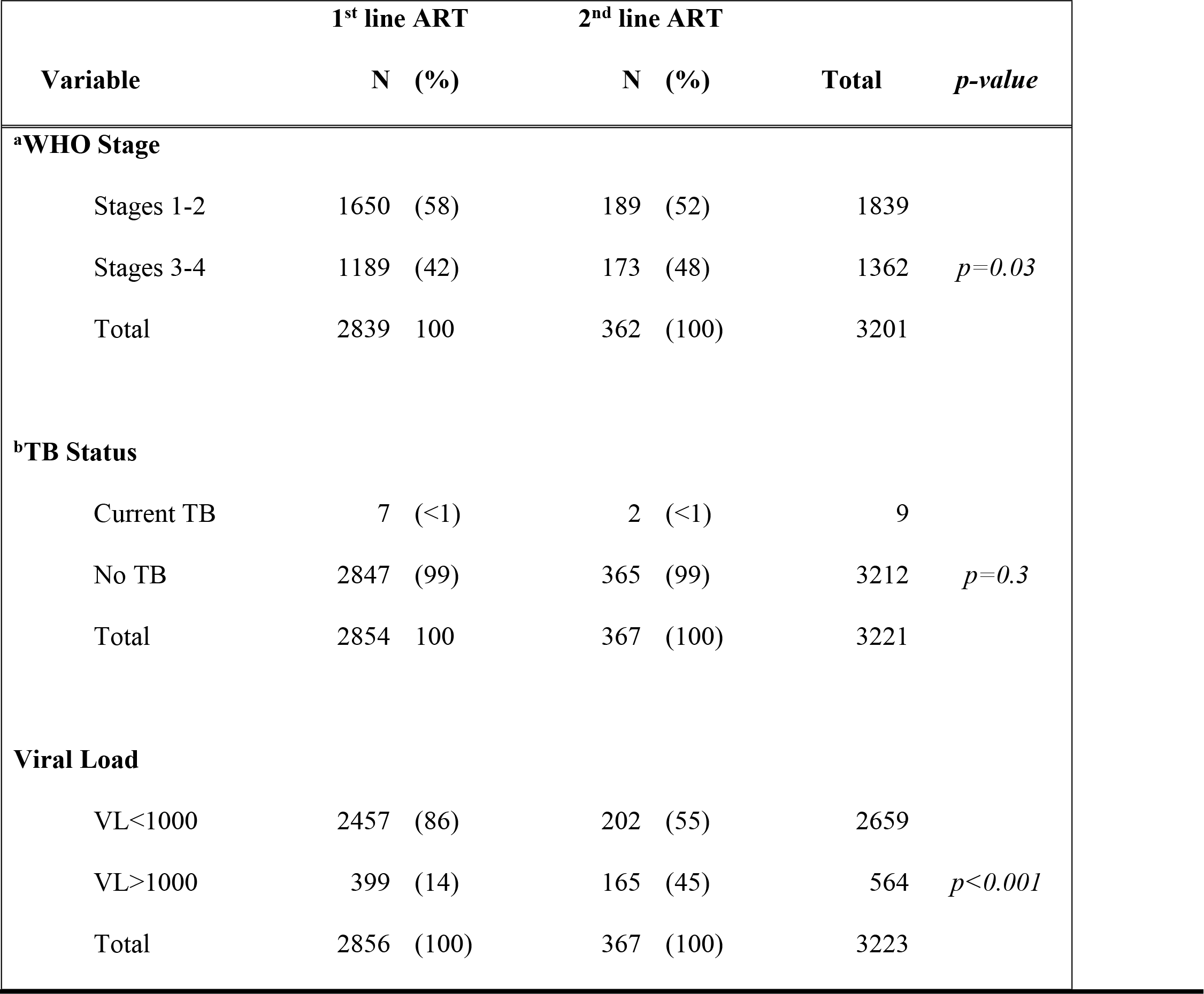
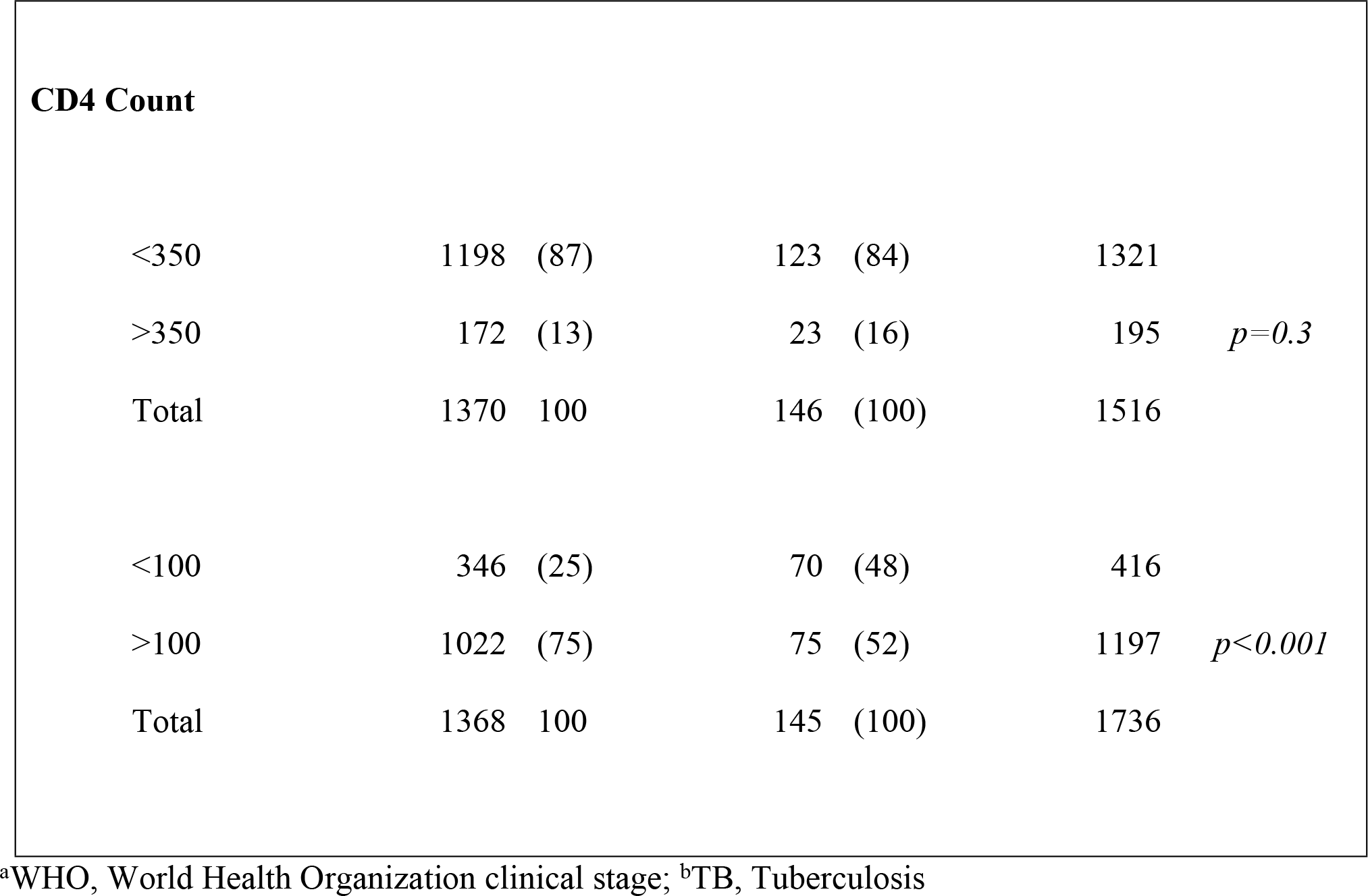
Clinical characteristics of HIV patients on 1^st^ and 2^nd^ line ART in Mutare District, Manicaland; Zimbabwe between 2015-2017 (N=3,223).

Log-binomial regression analysis was completed to determine the independent strength of risk factor association with viral load non-suppression among those on 1^st^ line ART, see **Table 3**. 1^st^ line ART patients had three times the risk of VL non-suppression if they were in the 0-9 years’ age group (adjusted RR=2.9; CI=1.7-4.8, *P*<0.001) and two times the risk if in the 10-19 years’ age group (adjusted RR=2.2, CI=1;4-3.2, *P*<0.001) compared to those 20-49 years. Additionally, 1^st^ line patients were at significantly higher risk of VL non-suppression if they had concurrent TB (adjusted RR=9.4; CI=3.0-29.7, *P*<0.001) and were male (adjusted RR=1.5; CI=1.1-2.1, *P*=0.02). There were no other significant associations among the 1^st^ line ART patients. A regression analysis completed among risk factors for viral load non-suppression for 2^nd^ line ART patients found no significant differences.

**Table 3.**
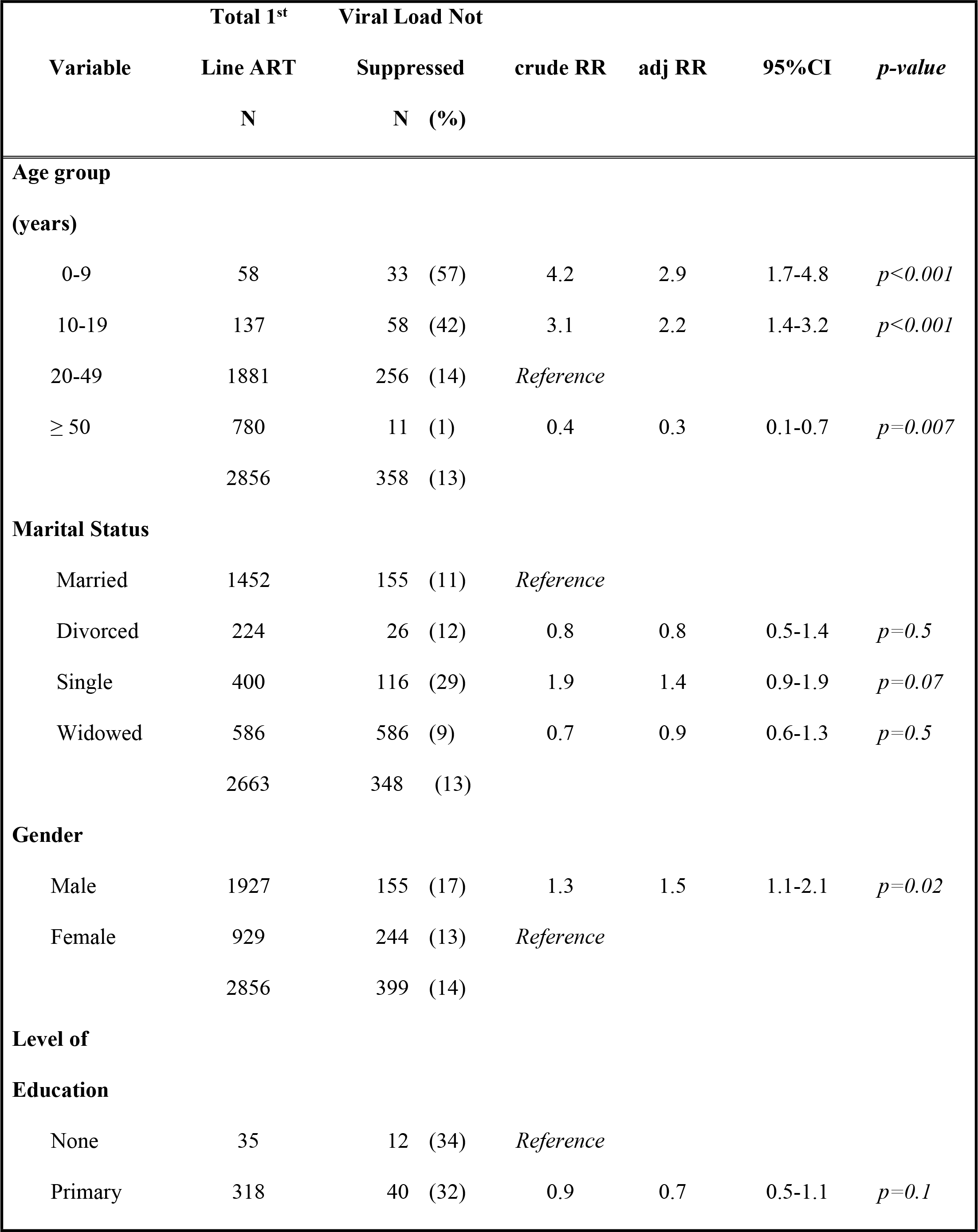
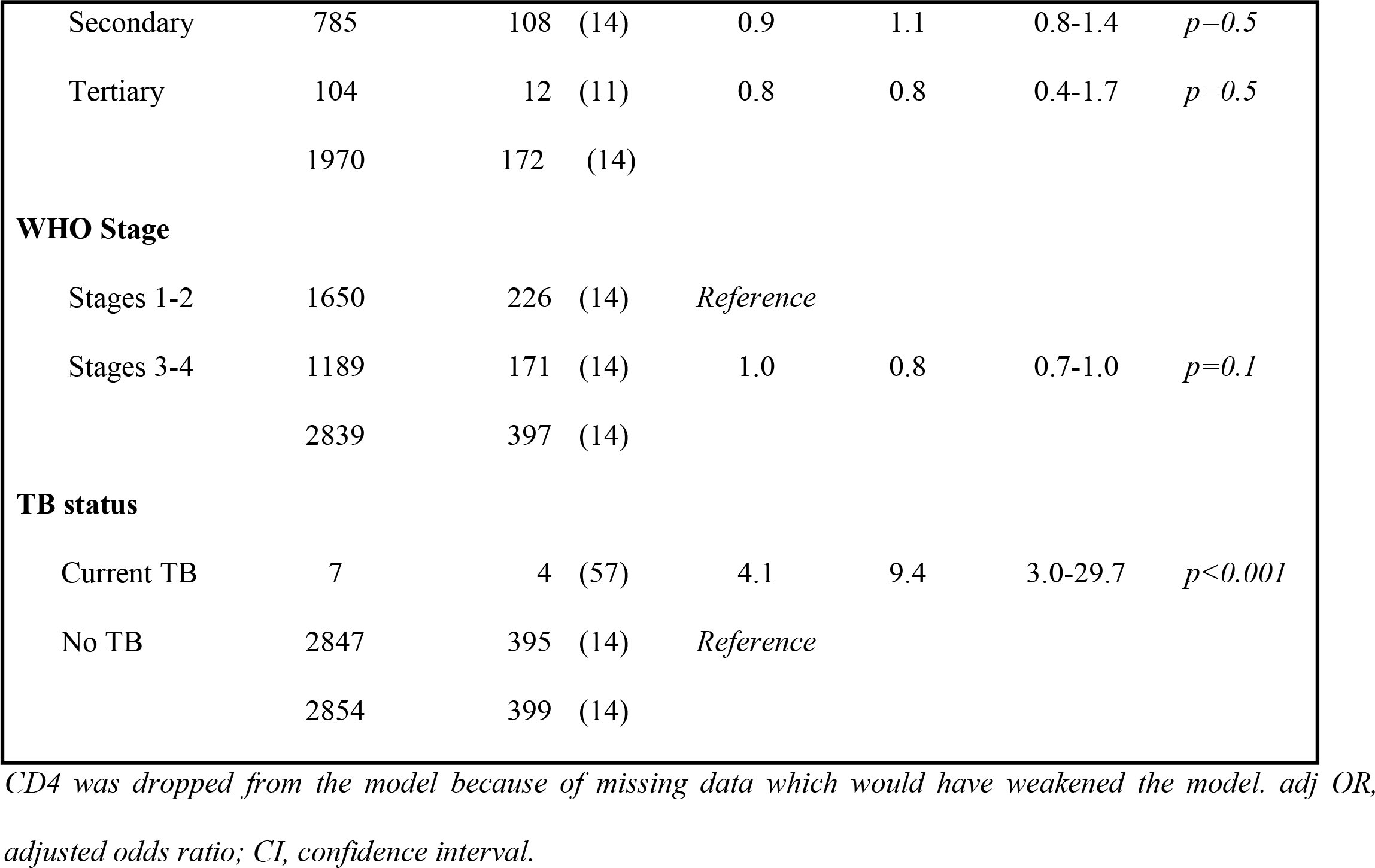
Multivariate analysis of factors associated with viral load suppression among HIV patients on 1^st^ line ART Mutare District, Manicaland; Zimbabwe 2015-2017 (N=2856).

|  | Total 1<sup>st</sup> | Viral Load Not |  |  |  |  |  |
| --- | --- | --- | --- | --- | --- | --- | --- |
| Variable | Line ART | Suppressed |  | crude RR | adj RR | 95%CI | *p-value* |
|  | N | N (%) |  |  |  |  |  |
| Age group |  |  |  |  |  |  |  |
| (years) |  |  |  |  |  |  |  |
| 0-9 | 58 | 33 (57) |  | 4.2 | 2.9 | 1.7-4.8 | *p<0.001* |
| 10-19 | 137 | 58 (42) |  | 3.1 | 2.2 | 1.4-3.2 | *p<0.001* |
| 20-49 | 1881 | 256 (14) |  | Reference |  |  |  |
| ≥ 50 | 780 | 11 (1) |  | 0.4 | 0.3 | 0.1-0.7 | *p=0.007* |
|  | 2856 | 358 (13) |  |  |  |  |  |
| Marital Status |  |  |  |  |  |  |  |
| Married | 1452 | 155 (11) |  | Reference |  |  |  |
| Divorced | 224 | 26 (12) |  | 0.8 | 0.8 | 0.5-1.4 | *p=0.5* |
| Single | 400 | 116 (29) |  | 1.9 | 1.4 | 0.9-1.9 | *p=0.07* |
| Widowed | 586 | 586 (9) |  | 0.7 | 0.9 | 0.6-1.3 | *p=0.5* |
|  | 2663 | 348 (13) |  |  |  |  |  |
| Gender |  |  |  |  |  |  |  |
| Male | 1927 | 155 (17) |  | 1.3 | 1.5 | 1.1-2.1 | *p=0.02* |
| Female | 929 | 244 (13) |  | Reference |  |  |  |
|  | 2856 | 399 (14) |  |  |  |  |  |
| Level of Education |  |  |  |  |  |  |  |
| None | 35 | 12 (34) |  | Reference |  |  |  |
| Primary | 318 | 40 (32) |  | 0.9 | 0.7 | 0.5-1.1 | *p=0.1* |

|  |  |  |  |  |  |  |
| --- | --- | --- | --- | --- | --- | --- |
| Secondary | 785 | 108 (14) | 0.9 | 1.1 | 0.8-1.4 | $p=0.5$ |
| Tertiary | 104 | 12 (11) | 0.8 | 0.8 | 0.4-1.7 | $p=0.5$ |
|  | 1970 | 172 (14) |  |  |  |  |
| **WHO Stage** |  |  |  |  |  |  |
| Stages 1-2 | 1650 | 226 (14) | *Reference* |  |  |  |
| Stages 3-4 | 1189 | 171 (14) | 1.0 | 0.8 | 0.7-1.0 | $p=0.1$ |
|  | 2839 | 397 (14) |  |  |  |  |
| **TB status** |  |  |  |  |  |  |
| Current TB | 7 | 4 (57) | 4.1 | 9.4 | 3.0-29.7 | $p<0.001$ |
| No TB | 2847 | 395 (14) | *Reference* |  |  |  |
|  | 2854 | 399 (14) |  |  |  |  |

CD4 was dropped from the model because of missing data which would have weakened the model. adj OR, adjusted odds ratio; CI, confidence interval.

## Discussion

Our study provides the first look at the most recent HIV outcomes since implementation and scale-up of routine viral load testing in Mutare district, Manicaland, Zimbabwe. The results reveal that there remains a significant amount of under-reporting and documentation of routine HIV VL monitoring in a district that is the capital of a province in Zimbabwe.

While the rate of reported VL suppression is close to the WHO 90% goal, at 86% for 1^st^ line ART patients, only 18% had reported VL results. Likewise, VL reporting was a challenge among 2^nd^ line ART patients, with only 35% having available VL results. The multivariate analysis revealed that those with unsuppressed VL among 1^st^ line patients were more likely to be < 20 years old, male, or have concurrent TB. The results suggest that increased programmatic focus on patients with these risk factors is needed to reduce the frequency of VL non-suppression.

The findings are unique because they provide “real-world” field perspective into the successes and challenges still facing a remote province in southern Zimbabwe and are likely comparable in other parts of the country and similar contexts elsewhere. The strengths of this study include the use of a district-wide implemented electronic patient monitoring tool for data sourcing and support from collaborative partners for VL sample transport, VL platform resources and medication supply.

The primary limitation is the lack of reported viral load results for both 1^st^ and 2^nd^ line ART patients which directly limits the ability to determine the true rate of VL suppression. There are likely several reasons for the lack of VL results being documented. First, there is a single VL platform based at Mutare Provincial Hospital (MPH). The MPH lab is the sole provider of all VL results for ART patients in both Manicaland (population 1.7 million) and Masvingo (population 1.5 million).(15) With a conservative estimate of 10% prevalence of HIV within these two provinces, that leaves approximately 300,000 patients needing at least yearly VL results provided by MPH, which is not technically feasible with current resources, regardless of partner support.

Second, VL sample transport has been reported to be a significant problem on several levels. (16) Mutare district is currently using a combination of whole blood or plasma VL samples for closer facilities and dried blood spot (DBS) samples from further distances. These samples need to be delivered to the MPH laboratory within specific time constraints, which is especially true for whole blood and plasma samples, requiring a secure cold-chain in place and can lead to increased rates of sample rejection(17)

Third, increased laboratory turnaround times have been previously reported as a limiting factor in reporting VL results in a timely fashion.(16) Fundamental to reaching the UNAIDS 90-90-90 HIV goals has been the scale-up and decentralization of VL testing platforms.(18) However, VL testing devices require significantly costly reagents that frequently require sponsored support, additional training and personnel to manage. Reagent stock-outs for VL testing remains challenging in many African contexts and maybe a key contributing factor at MPH(19) The MPH laboratory has limited human resources which can affect both the time to process incoming VL samples and VL result entry into the Laboratory Management Information System (LMIS). Lastly, increased turnaround times can paradoxically lead to reduced VL sample submission.(16)

The exact amount of VL access to those PLHIV on ART in Zimbabwe is currently uncertain. Previously, it has been reported that only 5.6% of ART patients had access to VL testing in 2015 in Zimbabwe.(19) Our study found that 18% of 1^st^ line patients had documented VL results, which demonstrates an improving trend, but far from the 90% target by the end of 2018 set by the Zimbabwe HIV Viral Load Scale-Up Plan 2015-2018.(4) More concerning is the increased risk of drug resistance mutations leading to HIV drug resistance that are more likely to occur in patients with undetected continued viremia secondary to the lack of VL testing(20)

Our study findings demonstrate near 90% VL suppression among those on 1^st^ line ART within the Mutare District of Manicaland. However, the lack of adequate VL testing coverage appears to be of significant concern on multiple levels. These results suggest that there is an urgent need for increased support for the Zimbabwe VL scale-up plan from collaborative partners which should likely target multiple components of testing system. Further study is needed determine exactly where the greatest gaps are in VL scale-up and our findings must serve as a call for others to assist in this effort. It is highly likely that these challenges in viral load monitoring are not isolated to Mutare district or to Zimbabwe alone.

## Conclusions

For those people living with HIV on 1^st^ line ART in Mutare district, Manicaland, Zimbabwe, the frequency of reported viral load results were found to be only 18% among those on 1^st^ line ART, while their rate of viral load suppression was near 90%. Viral load testing coverage appears to be lagging behind current Zimbabwe goals and increased support is needed to improve the quality of HIV care and help reduce the threat of possible HIV drug resistance in the future.

## Acknowledgement

This research was conducted through the Structured Operational Research and Training Initiative (SORT IT), a global partnership led by the Special Programme for Research and Training in Tropical Diseases at the World Health Organization (WHO/TDR). The training model is based on a course developed jointly by the International Union Against Tuberculosis and Lung Disease (The Union) and Medécins sans Frontières (MSF). The specific SORT IT program which resulted in this publication was implemented by the Centre for Operational Research, The Union, Paris, France. Mentorship and the coordination/facilitation of this particular SORT IT workshop was provided through the Centre for Operational Research, The Union, Paris, France; the Department of Tuberculosis and HIV, The Union, Paris, France; the University of Washington, School of Public Health, Department of Global Health, Seattle, Washington, USA; and AMPATH, Eldoret, Kenya.

## Supporting information

S1 Registered HIV Patients Mutare (XL)

S2 Mutare HIV patients 1^st^ line (XL)

S3 Mutare HIV patients 2^nd^ line (XL)

## References

1. UNAIDS (Joint United Nations Programme on HIV/AIDS). Ambitious treatment targets: writing the final chapter of the AIDS epidemic: writing the final chapter of the AIDS epidemic. 2014;1–36.

2. UNAIDS. Ending Aids Progress Towards the 90-90-90 Targets. Glob Aids Updat [Internet]. 2017;198. Available from: http://www.unaids.org/sites/default/files/media_asset/Global_AIDS_update_2017_en.pdf

3. Survey H. Zimbabwe. 2015;

4. National Viral Load Scale Up Plan 2015-2018.

5. Bulage L, Ssewanyana I, Nankabirwa V, Nsubuga F, Kihembo C, Pande G, et al. Factors Associated with Virological Non-suppression among HIV-Positive Patients on Antiretroviral Therapy in Uganda, August 2014-July 2015. BMC Infect Dis. 2017;17(1):326.

6. Jobanputra K, Parker LA, Azih C, Okello V, Maphalala G, Kershberger B, et al. Factors associated with virological failure and suppression after enhanced adherence counselling, in children, adolescents and adults on antiretroviral therapy for HIV in Swaziland. PLoS One. 2015; 10(2):1–12.

7. Costenaro P, Penazzato M, Lundin R, Rossi G, Massavon W, Patel D, et al. Predictors of Treatment Failure in HIV-Positive Children Receiving Combination Antiretroviral Therapy : Cohort Data From Mozambique and Uganda. 2018;4(1):39–48.

8. Davey DJ, Abrahams Z, Feinberg M, Prins M, Serrao C, Medeossi B, et al. Factors associated with recent unsuppressed viral load in HIV-1-infected patients in care on first-line antiretroviral therapy in South Africa. 2018;

9. Zimbabwe Ministry of Health and Child Care, PEPFAR, ICAP, Agency N. Zimbabwe Population-Based HIV Impact Assessment (ZIMPHIA) 2015-2016. 2016;(December 2016):1–4.

10. World Health Organisation. Zimbabwe: WHO statistical profile. World Heal Organ. 2015;1–3.

11. ZIMSTATS. Manicaland Zimstats pdf.

12. World Health Organization. Guidelines Guideline on When To Start Antiretroviral Therapy and on Pre-Exposure Prophylaxis for Hiv. World Heal Organ [Internet]. 2015;(September):78. Available from: http://www.who.int/hiv/pub/guidelines/earlyrelease-arv/en/

13. MSF IN ZIMBABWE ACTIVITY REPORT 2015. 2015;

14. National Medicines and Therapeutics Policy Advisory, Committee (NMTPAC), and, The AIDS and TB Directorate M of H and C, Care Z. Guidelines for Antiretroviral Therapy for the Prevention and Treatment of HIV in Zimbabwe. 2016;(December). Available from: https://aidsfree.usaid.gov/sites/default/files/zw_arv_therapy_prevention.pdf

15. Zimstat. Zimbabwe Population Census 2012. Popul Census Off [Internet]. 2012;(2012): 1–151. Available from: http://www.zimstat.co.zw/dmdocuments/Census/CensusResults2012/National_Report.pdf

16. Minchella PA, Chipungu G, Kim AA, Sarr A, Ali H, Mwenda R, et al. Specimen origin, type and testing laboratory are linked to longer turnaround times for HIV viral load testing in Malawi. PLoS One [Internet]. 2017;12(2):1–13. Available from: http://dx.doi.org/10.1371/journal.pone.0173009

17. Bonner K, Siemieniuk RA, Boozary A, Roberts T, Fajardo E, Cohn J. Expanding access to HIV viral load testing: A systematic review of RNA stability in EDTA tubes and PPT beyond current time and temperature thresholds. PLoS One. 2014;9(12): 1–13.

18. Block TM, Rawat S, Brosgart CL, Francisco S. HHS Public Access. 2017;216(Suppl 9):69–81.

19. Kilmarx PH, Simbi R. Progress and Challenges in Scaling Up Laboratory Monitoring of HIV Treatment. PLoS Med [Internet]. 2016;13(8):10–2. Available from: http://dx.doi.org/10.1371/journal.pmed.1002089

20. WHO, CDC, TheGlobalFund. Hiv Dru [Internet]. Hiv Drug Resistance Report 2017 Trends Quality Action. 2017. Available from: http://apps.who.int/iris/bitstream/10665/255896/1/9789241512831-eng.pdf?ua=1

